# Proof of concept: Molecular prediction of schizophrenia risk

**DOI:** 10.1101/129213

**Authors:** Anna R. Docherty, Arden Moscati, Daniel E. Adkins, Gemma T. Wallace, Guarav Kumar, Brien P. Riley, Aiden Corvin, F. Anthony O’Neill, Michael Gill, Kenneth S. Kendler, Patrick F. Sullivan, Ayman H. Fanous, Silviu-Alin Bacanu

**Affiliations:** Department of Psychiatry, University of Utah School of Medicine, Salt Lake City, UT USA; Department of Human Genetics, University of Utah School of Medicine, Salt Lake City, UT USA; Virginia Institute for Psychiatric and Behavioral Genetics, Virginia Commonwealth University School of Medicine, Richmond, VA USA; Department of Sociology, University of Utah, Salt Lake City, UT USA; Center for Biomarker Research and Personalized Medicine, Virginia Commonwealth University School of Medicine, Richmond, VA USA; Department of Psychiatry and Neuropsychiatric Genetics Research Group, Trinity Translational Medicine Institute, Trinity College Dublin, Dublin, Ireland; Queen’s University Belfast, Ireland; Departments of Genetics and Psychiatry, UNC-Chapel Hill, Chapel Hill, NC USA; Department of Medical Epidemiology and Biostatistics, Karolinska Institutet, Stockholm, Sweden; SUNY Downstate Medical Center, Brooklyn, NY USA; Washington Veterans Affairs Healthcare System, Washington, D.C. USA

## Abstract

**Key Points:** *Question:* To what extent do global polygenic risk scores (PRS), molecular pathway-specific PRS, complement component (C4) gene expression, MHC loci, sex, and ancestry jointly contribute to risk for schizophrenia-spectrum disorders (SZ)?

*Findings:* Global polygenic risk for schizophrenia, sex, and their interaction most robustly predict risk in a classification and regression tree model, with highest risk groups having 50/50 chance of SZ.

*Meaning:* Psychometric risk indicators, such as prodromal symptom assessments, may be enhanced by the examination of genetic risk metrics. Preliminary results suggest that of genetic risk metrics, global polygenic information has the most potential to significantly aide in the prediction of SZ.

**Abstract:** *Importance:* Schizophrenia (SZ) has a complex, heterogeneous symptom presentation with limited established associations between biological markers and illness onset. Many (gene) molecular pathways (MPs) are enriched for SZ signal, but it is still unclear how these MPs, global PRS, major histocompatibility complex (MHC) complement component (C4) gene expression, and MHC loci might jointly contribute to SZ and its clinical presentation. It is also unclear whether sex or ancestry interacts with these metrics to increase risk in certain individuals.

*Objective:* To examine multiple genetic metrics, sex, and their interactions as possible predictors of SZ risk. Genetic information could aid in the clinical prediction of risk, but it is still unclear which genetic metrics are most promising, and how sex interacts with genetic risk metrics.

*Design, Setting, and Participants:* To examine molecular risk in a proof-of-concept study, we used the Wellcome Trust case-control cohort and classified cases as a function of 1) polygenic risk score (PRS) for both whole genome and for 345 implicated molecular pathways, 2) predicted C4 expression, 3) SZ-relevant MHC loci, 4) sex, and 5) ancestry.

*Main Outcomes and Measures:* PRSs, C4 expression, SZ-relevant MHC loci, sex, and ancestry as joint risk factors for SZ.

*Results:* Recursive partitioning yielded 15 molecular risk classes and retained as significant psychosis classifiers only sex, genome-wide SZ polygenic risk, and one MP PRS. Sex was the most robust classifier in a stepwise regression, and there was a significant interaction of sex with SZ PRS on case status, suggesting males have a lower polygenic risk threshold. By down-sampling case proportion to 1% and 1.4% population base rates in males and females, respectively, high-risk subtypes defined by this model had roughly a 52% odds of developing SZ (individuals with SZ PRS elevated by 2.6 SDs; incidence = 51.8%).

*Conclusions and Relevance:* This proof-of-concept suggests that global SZ PRS, sex, and their interaction are robust predictors of risk and that males have a lower PRS threshold for onset. Implications for the integration of these metrics with psychometrically-identified risk are discussed.

## Introduction

Schizophrenia (SZ) has a complex, heterogeneous symptom presentation with limited established associations between biological markers and illness presentation. The Psychiatric Genomics Consortium (PGC) has made unprecedented advances in the molecular etiology of SZ^1,2^ by confirming that SZ is highly polygenic, with no aggregation of top loci effects accounting for more than 3% of the variance in case-control status.^3–6^ However, it has recently been recognized that for practical purposes, a genome-wide polygenic risk score (PRS) may be an informative genomic prediction metric.

Previous attempts to classify individuals based on genetic information have failed to account for sex, ancestry, major histocompatibility complex (MHC) factors, and molecular pathway information. It can be argued that all of these factors and their interactions could significantly contribute to relative risk. And given the enhanced cost-effectiveness of genotyping in recent years, it is important to consider how this information may complement the psychometric (interview and survey-based) tools we currently use to identify SZ risk.

For instance, *C4* gene expression^7^ in MHC regions and many molecular (gene) pathways^1^ (MPs) are associated with SZ. It is possible that these factors interact upon a background of elevated global SZ polygenic risk in the development of psychosis. It is of great interest to determine how global SZ PRS, polygenic risk specific to MPs, predicted *C4* expression, and sex may interact to give rise to incidence, relative risk, and symptom presentation. It is also possible that these factors may help reduce the genetic heterogeneity of SZ.

This proof of concept study reflects the development of a method for assessing the predictive utility of multiple genetic metrics in an SZ case-control sample. We evaluate the suitability of a method and of these implicated risk factors for clinical prediction. This method also serves to assign subjects to molecular subtypes based on the most robust genetic metrics. Considering the genetic complexity of SZ, the development of successful DNA-driven molecular classification of risk is highly desirable;^2^ in the context of precision medicine, identification of molecular subtypes predictive of onset, symptoms, functional outcome, or treatment response could lead to more effective interventions. Genetic classification may elucidate biological mechanisms of SZ, and the genetic overlap that we observe of SZ with other major psychiatric and medical disorders.^8–11^

This study used genome-wide association (GWAS) summary statistics from the largest available SZ cohort^1^ (N > 150,000) to construct subject-level genome-wide SZ PRS, as well as PRS specific to 345 human nominally enriched molecular pathways^1^, in data from the Wellcome Trust Case Control Consortium 2 (WTCCC2). Our approach differed from existing pathway enrichment analyses, based on summary statistics that regress out sex,^1^ because MPs may differ by sex. These data were then integrated with predicted *C4* expression and MHC single nucleotide polymorphism (SNP) data for *rs210133* and *rs1233604,*^7^ as well as data on sex and empirically-derived ancestry principal components (PCs). Affection status was then classified as a function of the above subject-level variables. This led to risk groups with a robust incidence elevation as a function of SZ PRS, sex, and SZ PRS*sex. A natural follow-up to this study will be to examine PGC cases and controls in a leave-one-out analysis.

## Methods

### Sampling

The SZ data were collected as part of the WTCCC2 study of 15 complex disorders and traits, and the dataset was made up of an Irish cohort of 3834 (1733 affected and 2101 unaffected) subjects.^12^ For all participants, DNA was extracted from whole blood using standard methods. Genomic DNA for all cases and control subjects was genotyped by the Sanger Institute, United Kingdom, or the Broad Institute, United States. Haplotype phasing and imputation was performed using IMPUTE2^13^ on samples that had passed quality control filtering. Imputation was performed using the HapMap2 and 3 and the 1000 Genomes reference panels.

### Molecular Pathway-Based and Genome-Wide Schizophrenia Polygenic Risk Scores

A list of implicated pathways was downloaded from the most recent PGC schizophrenia meta-analysis^1^ from Supplementary Table 5. Using R (v. 3.3.1),^14^ pathway names were matched to pathway database entries from GO (Gene ontology; http://www.geneontology.org), KEGG (Kyoto Encyclopedia of Genes and Genomes; http://www.genome.jp/kegg), PAN-PW (PANTHER; http://www.pantherdb.org/pathway), Reactome (http://www.reactome.org/download), BioCarta (downloaded from the Molecular Signatures Database v4.0 http://www.broadinstitute.org/gsea/msigdb/index.jsp), and NCI pathways (NCI: http://pid.nci.nih.gov). Gene lists corresponding to each pathway were then extracted from the relevant database with the number of genes per pathway ranging from 8 to 598 (mean = 9).

Next, using the biomaRt library Ensembl dataset within the Bioconductor R package^15^, the genes were matched to coordinate intervals using human assembly GRCh37 (the assembly relevant to the analysis by Ripke et al., 2014).^1^ All SNPs present in Ensembl for the coordinate intervals of the genes, ± 50 kb flanking regions^1^ were extracted and concatenated in non-redundant pathway specific SNP lists. Then strand direction and reference SNP coding were harmonized between testing and training data and ambiguous (i.e., A/T, G/C) SNPs were removed.

LDpred^16^ allows for the modeling of LD based on LD in the discovery sample to weigh the relative contributions of single variants to the outcome phenotype. LDpred uses postulated proportions of causal variants in the genome as Bayesian prior probabilities for GPS calculations, and a range of different priors were tested (proportions of 0.3, 0.1, 0.03, 0.01, 0.003, and 0.001, as well as the model of infinite variants of infinitesimally small effect) to construct scores. Scores were constructed for schizophrenia and for each biological pathway. The number of SNPs in each MP in these data can be found at: https://docs.google.com/spreadsheets/d/1mKcHwfVmprxJJLjrLDEgTQlrcf-7T86OZch0lBMZI5A. Prior to classification analyses, all pathway scores were LD-pruned and merged based on analyses of excessive correlation between scores (R^2^ < 0.5). While PGC2 coefficients were not estimated as leave-WTCCC2-out, our approach is not expected to be adversely affected due to i) WTCCC2 being a very small part of the overall sample and ii) the average-based nature of pathway scores, which tends to cancel out the (likely small) positive and negative overfitting biases associated with small *p-*values. This is supported by a large 0.98 estimated correlation between WTCCC2 and leave-WTCCC2-out scores.

### Major Histocompatibility Complex and Complement *C4* Data

We removed the extended MHC region (chr6:25-33Mb) from non-C4 related pathways because the single C4 locus and two SNPs included in these analyses account for a large portion of MHC region signals.^7^ In WTCCC2, the three most common C4 locus structures were present on multiple MHC SNP haplotypes. We used our 222 integrated haplotypes of MHC SNPs and C4 alleles as reference chromosomes for imputation. We found that the four most common structural forms of the C4A/C4B locus (BS; 7%, AL–BS; 31%, AL–BL; 41%, and AL–AL; 11%) could be inferred with reasonably high accuracy (0.7<r^2^<1). These 4 forms account for 90% of all haplotypes, so data loss from excluding others was minimal. SNP data were analyzed from 28,799 schizophrenia cases and 35,986 controls from 40 cohorts in 22 countries contributing to the Psychiatric Genomics Consortium. We evaluated association to 7,751 SNPs across the extended MHC locus (chr6: 25–34 Mb), to C4 structural alleles and to HLA sequence polymorphisms imputed from the SNP data. We also predicted levels of C4A and C4B expression from the imputed C4 structural alleles. Entered into the CART and general linear models were C4A, C4B, and C4A*C4B.

### Genetic Subgroup Analysis

Model-based recursive partitioning was used for the subgrouping analysis.^17^ Classification and regression trees (CARTs) were used to partition subjects by SZ PRS, biological pathway-based PRS, C4A, C4B, C4A/C4B and C4A*C4B gene expression, 10 empirically-derived ancestry PCs, and sex to predict case-control status. To construct CARTs that use only cuts that are statistically significant after adjusting for multiple testing of all possible cuts we used ctree function R party package.^18^ Ctree was used with a minimum cut criterion of 0.95 and Monte Carlo-based correction for multiple testing. After an optimal CART model was characterized, the selected CART variables and all their possible two-way interactions were entered into a step-wise regression model to find the best-fitting set of predictors.

## Results

The best fitting CART model is presented in Figure 1, and indicated very high incidence rates uniquely relating to global SZ PRS. Recursive partitioning yielded 15 molecular subtypes (CART leaves), and retained as significant classifiers sex, genome-wide SZ polygenic risk, and protein export pathway-based polygenic risk. Ancestry PCs were not significant classifiers of risk in this sample.

**Figure 1.**
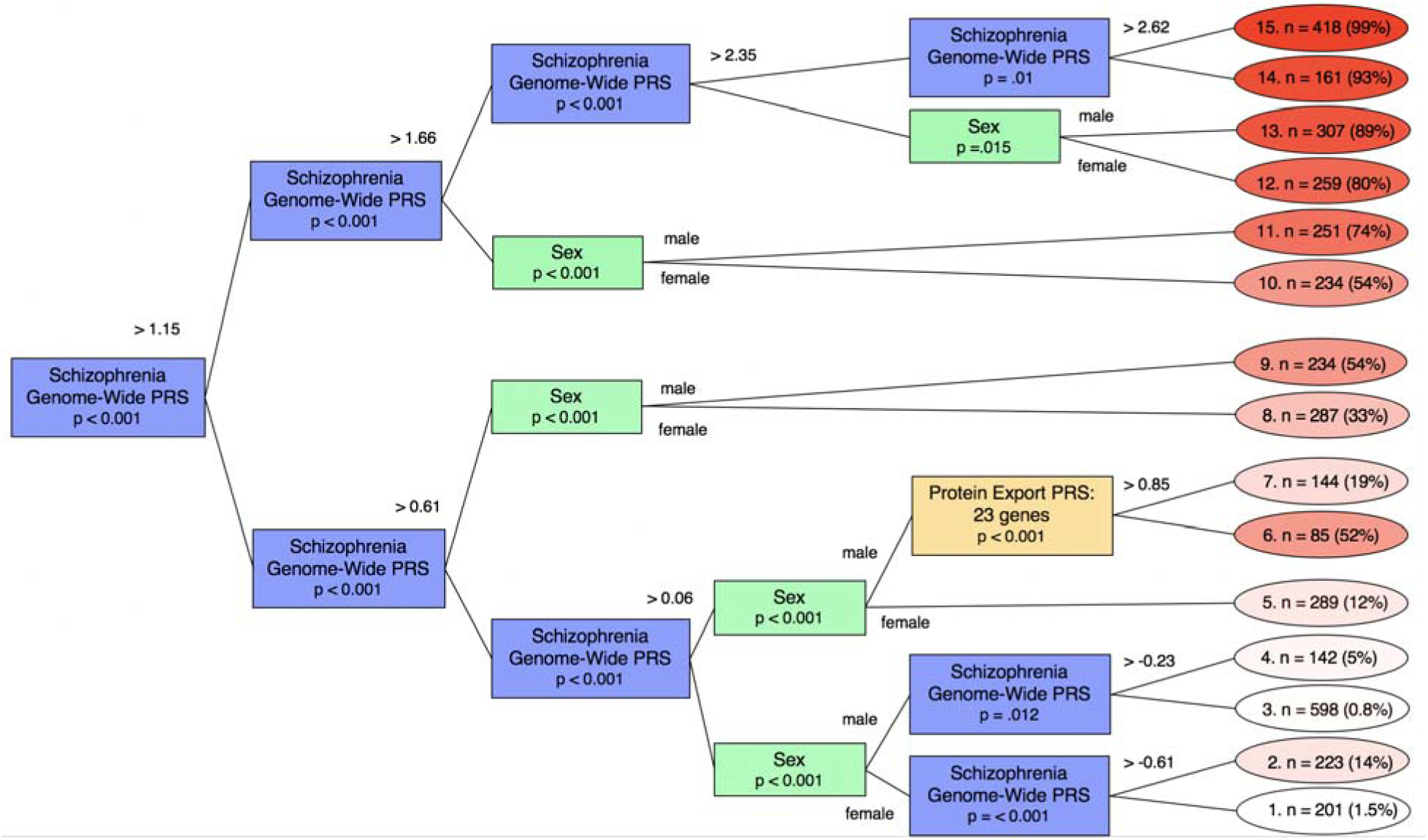
Classification tree model including SZ cases and controls. Presented here is a classification tree diagram featuring the partitioning factors, sex, and pathways that remained significant after multiple testing corrections. Identical nodes on different branches are colored similarly. The molecular risk classes are reflected in the leaves (ovals) with the number of individuals in the subtype, then % enrichment of cases. Red shading reflects the degree of enrichment, climbing to 99% in the group with highest molecular risk. Diagram paths between the nodes denote the PRS score at which the sample is split, or denote sex.

Stepwise regression statistics for the best fitting model are presented in Table 1. Sex was the most robust classifier in the stepwise regression, and the protein export pathway was excluded from the best-fitting model. Of note, there was a significant sex*PRS interaction, and the stepwise model captured a significant increase in male risk by genome-wide SZ PRS level (Figure 2). Incidence for males was higher than for females as a function of the same SZ PRS. PRSs from molecular pathways, and data on predicted *C4* expression, MHC SNPs, ancestry components, and all two-way interactions were not retained by any general linear model.

**Table 1.** Best Fitting Stepwise Regression Model with Schizophrenia PRS, Molecular Pathway PRSs, 10 Ancestry Principal Components, Sex, C4 Expression, and All 2-Way Interactions *(N = 3833)*

|  | **Estimate** | **SE** | **Z** | **p-value** |
| --- | --- | --- | --- | --- |
| Global SZ PRS | 1.33 | 0.19 | 7.01 | $2.36 \times 10^{-12}$ |
| Sex | -1.38 | 0.16 | -8.47 | $2.53 \times 10^{-17}$ |
| Global SZ PRS*Sex | 0.42 | 0.13 | 3.34 | $8.45 \times 10^{-4}$ |

*Note:* All components of the best-fitting model had significant p-values after correction for multiple testing. SZ = schizophrenia.

**Figure 2.**
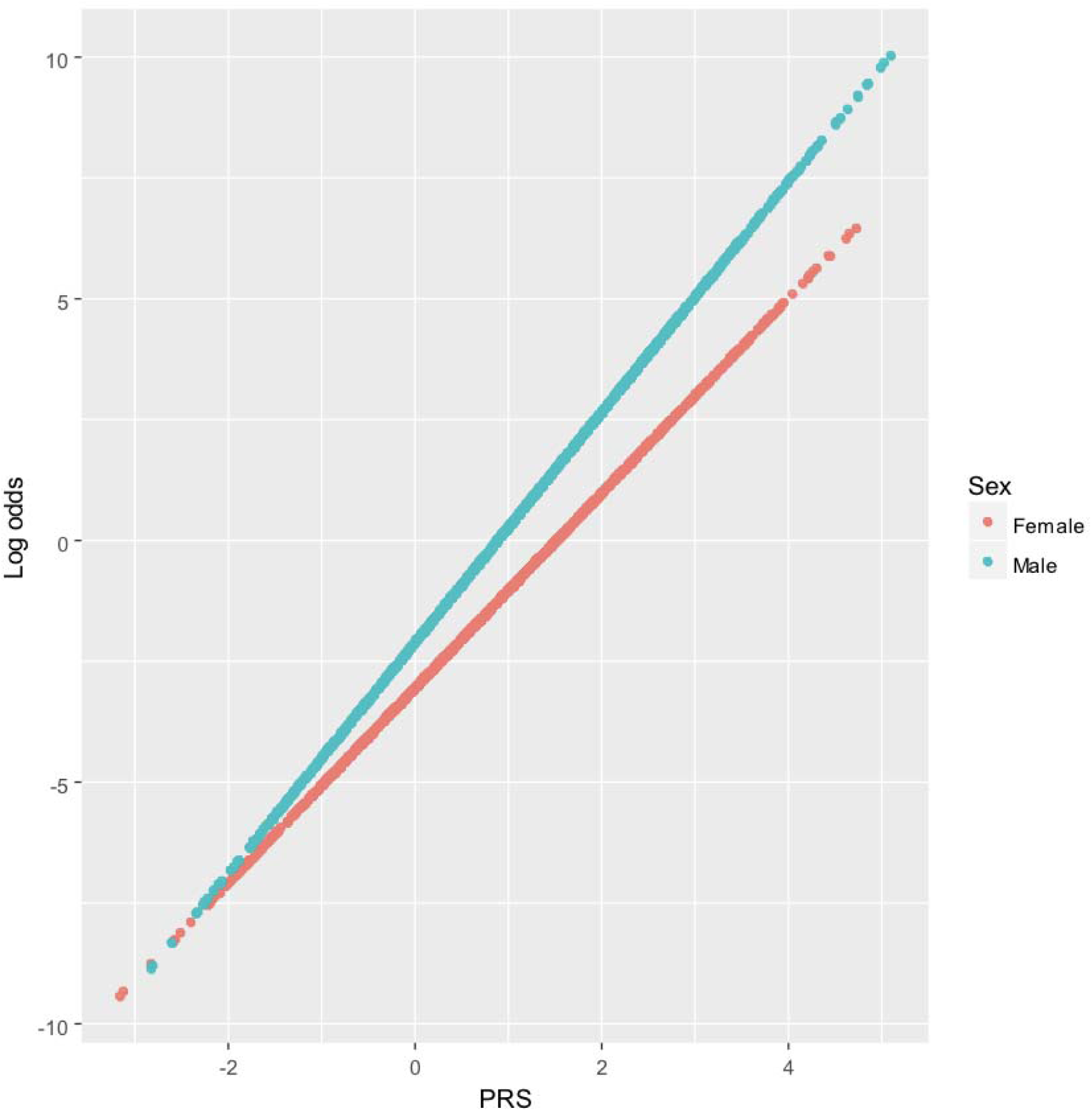
Log odds of being a case as a function of global SZ PRS and sex. As presented in Table 1, the best fitting model to predict case status included sex and the interaction of PRS with sex.

SZ incidence rates and relative risk are presented in Table 2. By down-sampling case proportion to 1% in females and 1.4% in males (expected population base rates), the high-risk subtypes defined by this model had approximately a 50/50 chance of developing SZ. These individuals had SZ PRS > 2.6 SDs from the mean of controls (incidence = 51.8%). This means that a standardized SZ PRS score >2.6 indicates a probability of psychosis of similar magnitude to being the monozygotic twin of someone with schizophrenia.

**Table 2.** Molecular Subtypes: Psychosis Relative Risk and Incidence Rates

| Subtype | Cases (%) | RR | Incidence (%) |
| --- | --- | --- | --- |
| 1 | 1.5 | 0.0 | 0.0 |
| 2 | 13.9 | 0.2 | 0.2 |
| 3 | 0.8 | 0.0 | 0.0 |
| 4 | 4.9 | 0.1 | 0.1 |
| 5 | 11.8 | 0.1 | 0.1 |
| 6 | 51.8 | 1.5 | 1.1 |
| 7 | 19.4 | 0.3 | 0.2 |
| 8 | 32.8 | 0.5 | 0.5 |
| 9 | 54.3 | 1.7 | 1.2 |
| 10 | 54.3 | 1.2 | 1.2 |
| 11 | 74.9 | 4.1 | 2.9 |
| 12 | 80.3 | 4.0 | 4.0 |
| 13 | 88.6 | 9.9 | 7.1 |
| 14 | 93.2 | 14.0 | 11.8 |
| 15 | 99.3 | 63.0 | 51.8 |

*Note:* Subtype = the number corresponding to each molecular subtype in Figure 1. Cases refers to the proportion of SZ cases in each subtype. RR = relative risk. Incidence refers to the predicted population incidence. The prevalence of schizophrenia is 1 and 1.4% in females and males, respectively.

## Conclusion

Current predictive risk indicators for SZ may be enhanced by using genetic risk metrics. Based on the preliminary results here, genetic information can aide in the prediction of who should be targeted in early prevention efforts. While these data are slightly more challenging to obtain than data from high-risk questionnaires and interviews, they are relatively cost-effective, easy to ascertain, and rely on the measurement of common variants. How psychometric prediction could improve with the integration of genetic risk metrics remains to be seen, and should be studied.

This proof-of-concept suggests that optimal clinical information for future prediction studies should include both global SZ PRS and sex. Though these findings need independent replication, individuals in the highest PRS group have > 50% SZ incidence, and when PRS drops to below 2.3 SDs from the control mean, the risk of psychosis drops significantly.

The observation of a PRS by sex interaction is notable, and suggests that for individuals above a certain SZ PRS threshold, being male elevates risk compared to being female. This is one explanation of higher base rates in males in the general population, but these results must be replicated in larger cohorts.

There are some important limitations to the current analysis. First, the sample is genetically homogeneous given that participants were derived from a Northern European population. Thus, effects of ancestry on risk cannot be sufficiently studied in this sample. Future research will need to replicate these effects in samples of other ancestry. Second, the WTCCC2 sample is a small subset of the larger PGC2 meta-analysis from which discovery summary statistics were derived. This could have biased the predictive results in favor of polygenic risk metrics. However, given that WTCCC2 was just a small fraction of the overall PGC2 sample, while being adequately powered, that bias is minimal. Third, and importantly, this sample serves only to illustrate our proof of concept; again, results therefore require replication in larger cohorts.

It is possible that high-risk PRS groups will be phenotypically distinct in samples with deeper phenotyping data and adequate power to detect effects on symptoms and course. Future research would also benefit from studies of the correspondence between SZ polygenic risk metrics and prodromal or high-risk traits. It is our hope that an application of these methods to larger and more diverse samples, with deeper phenotyping, will yield additional meaningful insights to improve risk prediction and targeted early intervention.

## Cited References

1. Ripke S, Neale BM, Corvin A, et al. Biological insights from 108 schizophrenia-associated geneti loci. Nature. 2014;511(7510):421–427.

2. Jablensky A. Schizophrenia or schizophrenias? The challenge of genetic parsing of a complex disorder. Am J Psychiatry. 2015;172(2):105–107.

3. Cross-Disorder Group of the Psychiatric Genomics Consortium. Identification of risk loci with shared effects on five major psychiatric disorders: a genome-wide analysis. The Lancet. 2013;381(9875):1371–1379.

4. Kendler KS, O’Donovan MC. A breakthrough in schizophrenia genetics. Jama Psychiat. 2014;71(12):1319–1320.

5. Sivakumaran S, Agakov F, Theodoratou E, et al. Abundant Pleiotropy in human complex diseases and traits. Am J Hum Genet. 2011;89(5):607–618.

6. Sullivan PF, Daly MJ, O’Donovan M. Genetic architectures of psychiatric disorders: the emerging picture and its implications. Nat Rev Genet. 2012;13(8):537–551.

7. Sekar A, Bialas AR, de Rivera H, et al. Schizophrenia risk from complex variation of complement component 4. Nature. 2016;530(7589):177–183.

8. Bulik-Sullivan B, Finucane HK, Anttila V, et al. An atlas of genetic correlations across human diseases and traits. Nat Genet. 2015;47(11):1236–1241.

9. Cross-Disorder Group of the Psychiatric Genomics Consortium. Genetic relationship between five psychiatric disorders estimated from genome-wide SNPs. Nat Genet. 2013;45(9):984–995.

10. Purcell SM, Wray NR, Stone JL, et al. Common polygenic variation contributes to risk of schizophrenia and bipolar disorder. Nature. 2009;460(7256):748–752.

11. Sullivan PF, Magnusson C, Reichenberg A, et al. Family history of schizophrenia and bipolar disorder as risk factors for autism. Arch Gen Psychiat. 2012;69(11):1099–1103.

12. Irish Schizophrenia Genomics Consortium, the Wellcome Trust Case Control Consortium. Genome-wide association study implicates HLA-C*01:02 as a risk factor at the major histocompatibility complex locus in schizophrenia. Biol Psychiatry. 2012;72(8):620–628.

13. Howie BN, Donnelly P, Marchini J. A flexible and accurate genotype imputation method for the next generation of genome-wide association studies. Plos Genet. 2009;5(6):e1000529.

14. R Core Team. R Foundation for Statistical Computing. 2014; http://www.r-project.org/

15. Durinck S, Spellman PT, Birney E, Huber W. Mapping identifiers for the integration of genomic datasets with the R/Bioconductor package biomaRt. Nat Protoc. 2009;4(8):1184–1191.

16. Vilhjalmsson BJ, Yang J, Finucane HK, et al. Modeling linkage disequilibrium increases accuracy of polygenic risk scores. Am J Hum Genet. 2015;97(4):576–592.

17. Zeileis A, Hothorn T, Hornik K. Model-based recursive partitioning. J Comput Graph Stat. 2008;17(2):492–514.

18. Hothorn T, Hornik K, Zeileis A. Unbiased Recursive Partitioning: A Conditional Inference Framework. J Comput Graph Stat. 2006;15(3), 651–674.

